# Hybrid protein-ligand binding residue prediction with protein language models: Does the structure matter?

**DOI:** 10.1101/2023.08.11.553028

**Authors:** Hamza Gamouh, Marian Novotný, David Hoksza

## Abstract

**Background:** Predicting protein-ligand binding sites is crucial in studying protein interactions with applications in biotechnology and drug discovery. Two distinct paradigms have emerged for this purpose: sequence-based methods, which leverage protein sequence information, and structure-based methods, which rely on the three-dimensional (3D) structure of the protein. We propose to study a hybrid approach combining both paradigms’ strengths by integrating two recent deep learning architectures: protein language models (pLMs) from the sequence-based paradigm and Graph Neural Networks (GNNs) from the structure-based paradigm. Specifically, we construct a residue-level Graph Attention Network (GAT) model based on the protein’s 3D structure that uses pre-trained pLM embeddings as node features. This integration enables us to study the interplay between the sequential information encoded in the protein sequence and the spatial relationships within the protein structure on the model’s performance.

**Results:** By exploiting a benchmark dataset over a range of ligands and ligand types, we have shown that using the structure information consistently enhances the predictive power of baselines in absolute terms. Nevertheless, as more complex pLMs are employed to represent node features, the relative impact of the structure information represented by the GNN architecture diminishes.

**Conclusions:** The above observations suggest that, although using the experimental protein structure almost always improves the accuracy binding site prediction, complex pLMs still contain structural information that lead to good predictive performance even without using 3D structure.

## 1 Introduction

Proteins are fundamental biomolecules that play a critical role in the functioning of all living organisms. They are involved in various biological processes such as signal transduction or cell regulation and interact with other macromolecules and small molecules to perform their functions. The interaction is mediated through binding sites on the protein surface. These binding sites contain residues crucial for the ligand molecule’s recognition and binding. Thus, the study of protein-ligand binding sites and binding residues is essential for understanding the fundamental mechanisms of biological processes with profound impact on applications such as drug discovery [1, 2] and biotechnology [3].

With the rapid advances in computational techniques in the last two decades, various methods have been developed for detecting protein-ligand binding sites. The methods use diverse algorithms and exploit different types of information from protein sequences and 3D structure, broadly categorizing the approaches into sequence-based and structure-based methods [4, 5].

Before describing the existing methods, we should emphasize that the problem of predicting protein-ligand interactions can be approached in two main ways: binding residue prediction, where sequence-based methods are mainly used, and binding site prediction, where structure-based methods are the most appropriate. Binding residue prediction involves labeling individual residues of the protein depending on whether they belong to a binding site. In contrast, binding site prediction aims at detecting surface regions capable of accommodating ligands that can potentially bind to the protein.

Sequence-based methods operate on amino acid sequences and are characterized by their ability to identify binding residues solely from protein sequence data. Although sequence-based methods can only predict individual binding residues and not full binding sites they can still be relevant in many applications, such as variant effect prediction (VEP) as the mutation of a binding residue increases the probability of a detrimental impact of such mutation by hampering the protein’s ability to bind ligands [3].

Traditional sequence-based tools, such as ConSurf [6] and S-Site [7], are template-based methods that use proteins with known binding sites as templates together with the evolutionary conservation information to predict binding residues from highly conserved regions of the protein.

In contrast, more recent methods rely on machine learning algorithms to make predictions. With the exponential increase in the size of biological databases [8], there has been an explosion of machine learning methods to solve all kinds of tasks in bioinformatics [9]. In the context of sequence-based methods for protein-ligand binding site prediction, different machine learning-based methods utilize different types of information about a protein sequence and its amino acids.

Several methods use Support Vector Machines (SVM) and Random Forest (RF) as their main classification algorithms and various input features. TargetS [10] constructs features using evolutionary information from Position Specific Scoring Matrix (PSSM), predicted secondary structure, and ligand-specific binding propensities of residues. ATPint [11] utilizes evolutionary information, hydrophobicity, and other predicted features such as average accessible surface area. NsitePred [12] computes features from the predicted secondary structure and uses additional information such as the predicted relative solvent accessibility (RSA) and dihedral angles, as well as PSSM features and residue conservation scores. LigandDSES [13] and LigandRFs [14] use amino acid physico-chemcial properties provided by the AAIndex database [15].

Deep learning methods have attracted enormous attention of bioinformaticians in recent years [16] due to their potential of automatic learning of complex representations from vast amounts of available data and due to their recent success in other fields, such as Natural Language Processing (NLP) [17] and Computer Vision (CV) [18]. Deep learning has also been used for binding residue detection in methods such as Deep-Bind [19] and DeepCSeqSite [20]. These approaches use Convolutional Neural Networks (CNNs) on protein sequences to predict binding residues. DeepBind uses residue types as input features, while DeepCSeqSite relies on various types of information, such as position-specific scoring matrix (PSSM), secondary structure (SS), dihedral angle (DA), and conservation scores (CS).

Recently, language models (LMs) have emerged as a viable option to represent protein sequences. Large LMs have become the standard method in NLP [21] due to their remarkable performance in a wide range of language-related tasks. An example of a very successful LM is the famous ChatGPT, based on the GPT-3 architecture [22], which can generate human-like responses in conversation. In bioinformatics, LMs have also been applied to address various challenges related to protein analysis [23–25].

A LM is a deep learning model architecture that is trained to learn complex representations of text input, also called embeddings, from an extensive corpus of text. LMs are built upon two basic successful ideas in NLP: masked language modeling and Transformer architecture. Masked language modeling [26] is a self-supervised learning strategy based on masking parts of the text and training the model to predict the missing parts. This strategy benefits from vast amounts of available unannotated data and forces the model to learn general embeddings that can be fine-tuned on downstream tasks where the data is scarce. The Transformer architecture [27] relies on the famous attention mechanism that helps the model attend only to relevant parts of the input by learning the attention weights of different parts of a text input.

Treating protein amino acids as words and sequences as sentences of a natural language opens a way to apply language modeling techniques to proteomics. Recently, several protein language models (pLMs) [28] were constructed by training Transformer architectures on large protein sequence datasets. The learned embeddings of protein sequences were then successfully applied to the prediction of various protein characteristics, such as protein structure [29, 30], or protein-protein interactions [31, 32]. In our recent work, we explored the potential of pLMs to predict protein-ligand binding residues showing superior performance over several state-of-the-art methods on multiple datasets [33]. In a broader view, the binding residue prediction problem can be viewed as a type of more general task of protein residue annotation, such as posttranslational modification prediction, where, indeed, pLMs have also been successfully applied [34, 35].

On the other hand, structure-based methods for protein-ligand binding site prediction utilize features derived from the protein 3D structure. Different structure-based methods vary in the way of representing the 3D protein structure and in the algorithm used for making the predictions.

FINDSITE [36] is a 3D template-based method that uses a threading algorithm based on binding-site similarity to groups of template structures. 3DLigandSite [37] and FunFOLD [38] are also template-based methods that combine sequence and structure similarity to extract homologous proteins from PDB from which ligands are extracted, superimposed, and clustered to determine the binding site associated with each cluster. Various other methods apply geometrical measurements over the 3D structure to detect cavities or hollows on the protein’s surface. SURFNET [39] is a method that positions spheres within the space between two protein atoms. LIGSITE [40] detects pockets with a series of simple operations on a cubic grid. FPocket [41] is based on Voronoi tessellation and alpha spheres. CurPocket [42] defines the binding sites by identifying clusters of concave regions from the curvature distribution of the protein surface. Methods such as Q-SiteFinder [43], FTSite [44] and SiteComp [45] are energy-based methods. Such methods place probes on the protein surface and subsequently locate cavities by estimating the energy potentials between the probes and the cavities. In addition to template-based, geometry-based, and energy-based methods, machine learning methods rely on 3D structural features, sometimes combined with other features, to train various machine learning algorithms. For instance, P2Rank [46] labels solvent-accessible surface points of the protein by using the Random Forest algorithm on a set of handcrafted physicochemical and structural features. The ligandable points are then clustered to obtain the binding pockets. Recently, deep-learning methods have been introduced for structure-based binding residue/site prediction as well. Often, the methods represent the protein structure as a 3D grid of voxels and use a 3D Convolutional Neural Network (CNN) [47] as their primary model architecture to learn the binding sites. These methods differ mainly in the input features and model hyperparameters. DeepSite [48], PUResNet [49] and DeepSurf [50] employ atomic chemical properties, DeepDrug3D [51] is based on interaction energies of ligand atoms with protein residues, while Deeppocket [52] uses atom types. More recent methods such as SiteRadar [53], GraphPLBR [54], EquiPocket [55], GraphBind [56] and GraphSite [57] use different variations of the Graph Neural Network (GNN) architecture and have demonstrated state-of-the-art performance.

GNN is a class of neural networks designed to operate on graphs and other structured data [58]. GNNs are based on the idea of representing the input data as a graph and propagating node information between the graph nodes. Each node is associated with a feature vector containing the node features. These features are iteratively updated by aggregating information from neighboring nodes using a series of message-passing steps. This property of GNNs enables the model to capture the graph’s local structure and learn more structure-based and context-aware embeddings. Methods based on GNNs may also benefit from large libraries of predicted protein structures by methods like AlphaFold [59, 60]. The primary output of a GNN is node feature vectors, which can be used for various node-level and graph-level downstream tasks. In recent years GNNs have been applied extensively in bioinformatics and have shown state-of-the-art results across multiple tasks [61].

In the following sections, we analyze the interplay of protein sequence and structure information by building a machine learning model that exploits two recent state-ofthe-art deep learning architectures; a Graph Neural Network augmented with protein-language model embeddings. Particularly, we want to address the following research questions: Can we improve the prediction performance by fusing both approaches? How much does the structure information from GNNs contribute to the predictive power of the solely sequence-based pLMs?

## 2 Methods

The high-level view of our approach, sketched in figure 1, is as follows. The first input of the pipeline is the protein sequence of single-letter amino acid codes. The sequence is processed by a pLM (Embedder), which computes embeddings of each amino acid in the sequence, i.e., residue-level embeddings. The second input is the corresponding protein 3D structure, described as a set of atom 3D coordinates. The structure is converted to a graph by the protein graph constructor (described in ***Protein graph construction***). In the protein graph, nodes correspond to residues labeled by the residue-level embeddings and edges to residues close in the 3D space. The protein graph is then processed by a GNN that predicts binding probabilities for each residue. Using a threshold, the predicted probabilities are converted to binding residue labels (binding vs. non-binding).

**Fig. 1:**
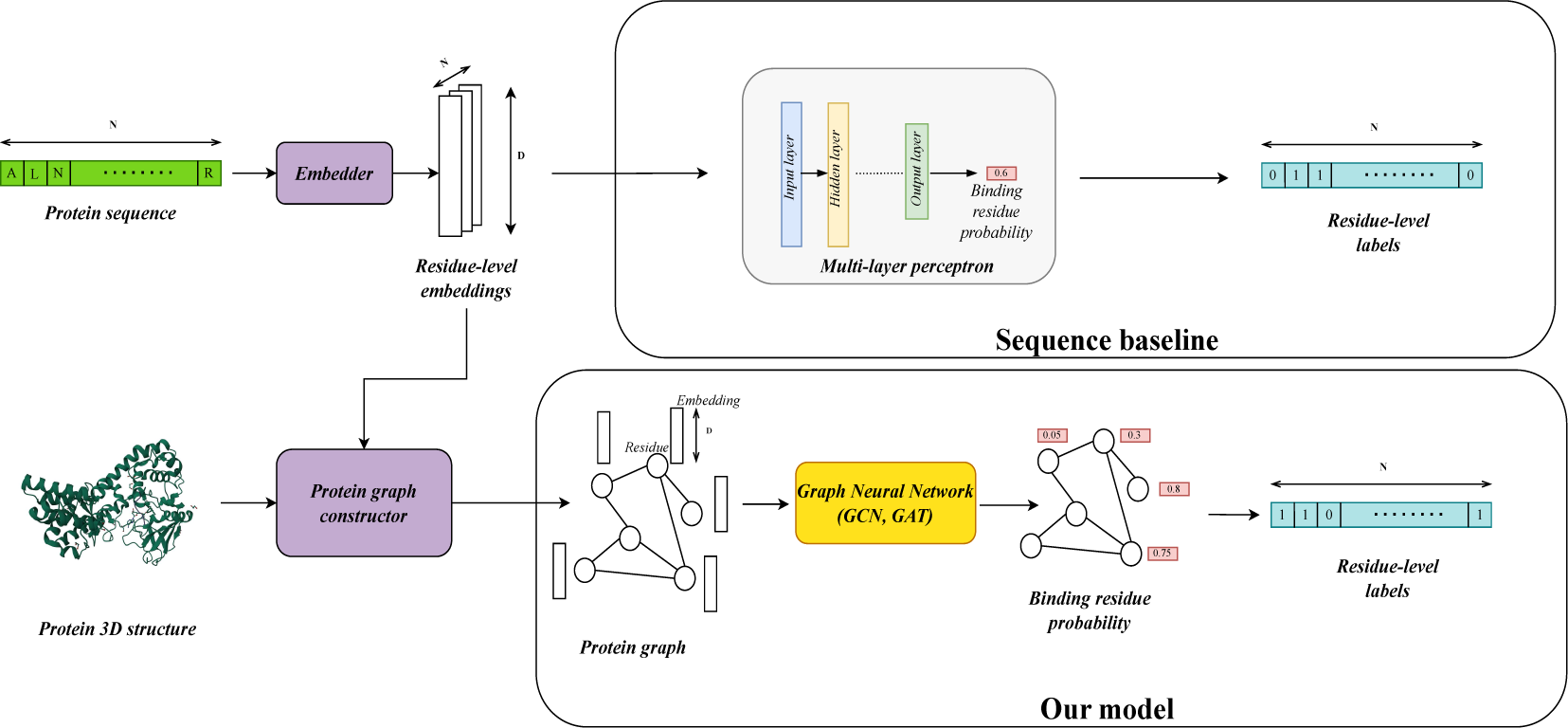
General architecture of our models

Furthermore, we measured the effect of the structure information that comes from the GNN models by comparing them to a baseline model, which is a sequence-based model that lacks graph structure information. The sequence baseline takes the residue-level embeddings as input and feeds it to a multi-layer perceptron which predicts the binding residue probability.

As mentioned, we use the Graph Neural Network (GNN) as our primary model architecture. Different GNN architectures vary in how they aggregate information from other nodes to transform the feature vectors. In our approach, we compare two well-known GNN architectures - Graph Convolutional Network (GCN) [62] and Graph Attention Network (GAT) [63].

The GCN uses convolutional operations to learn feature representations of nodes in a graph. The principle of GCNs is based on the idea of adapting convolutional neural networks (CNNs) [47] to the graph domain by replacing the regular grid-like structure of image data with an irregular graph structure. By analogy, GCNs define a convolution operation on graphs, which involves aggregating information from the node’s neighbors and updating the node’s feature representation accordingly. The graph convolution works by learning a trainable weight matrix shared across all nodes enabling the GCN to learn a set of filters specific to the graph structure.

The GAT follows the trend of the attention mechanism of the NLP Transformer architectures [27]. The model attends differently to different parts of a given node neighborhood by assigning importance scores to each neighbor based on their relevance to the current node. The attention mechanism enables the GAT to focus on the most relevant nodes in the graph while ignoring noise and irrelevant information. Figure 2 shows the architectural differences between GCNs and GATs.

**Fig. 2:**
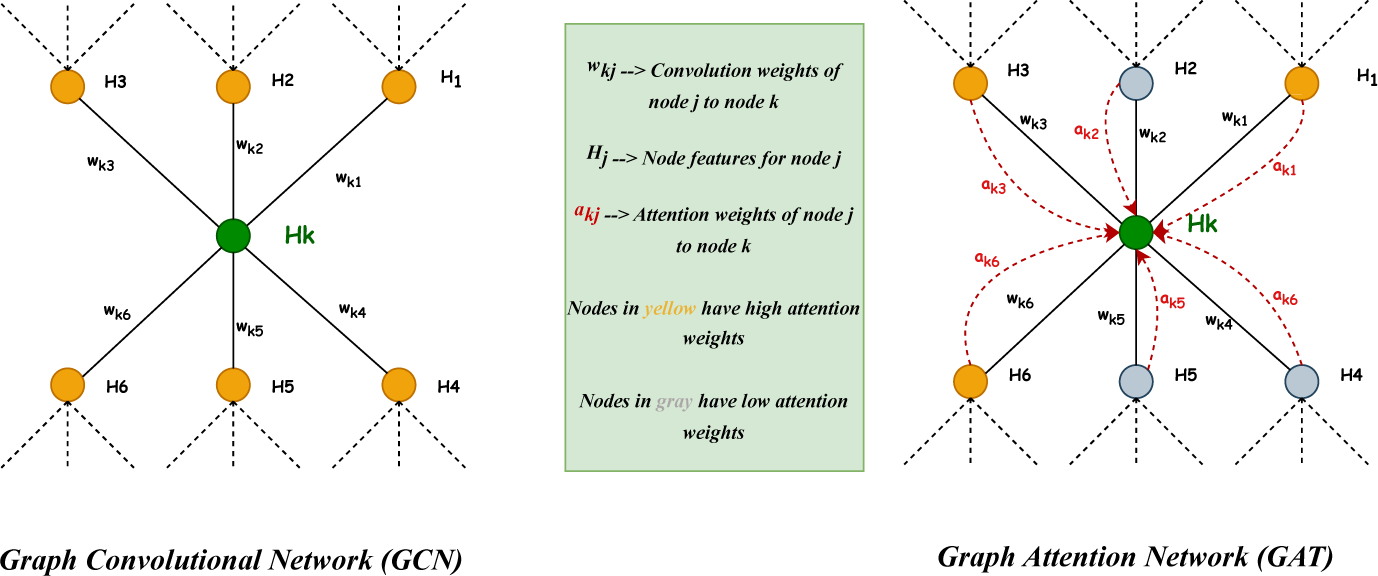
**Comparison of GCN and GAT architectures**

### 2.1 Protein graph construction

To use the GNN architecture, the protein needs to be represented as a graph with node features. In general, the strength of electrostatic interactions is inversely proportional to the distances between atoms. Therefore, it is physically plausible to enable information sharing between parts of the protein close to each other. Therefore, to construct the protein graph, we started with the 3D structure of the protein, and we constructed a proximity graph on the residue level. Nodes correspond to residues of the protein, and edges represent the closeness relationship of the residues to each other. Two residues are connected if the distance between their alpha-carbon atoms is less than the threshold distance. In this work, we explored the following thresholds: 4, 6, 8, and 10 ^°^A.

### 2.2 Protein language model embeddings

PLMs process sequences of amino acid letters and return two kinds of embeddings: an embedding for the whole protein sequence and an embedding for each sequence letter, i.e., residue-level embeddings. The latter embeddings can be directly used as node features of the protein graph. In this work, we used the following pLMs : two pLMs that are part of the ProtTrans project [64], ProtBERT-BFD that was pre-trained on BFD [65], and ProtT5-XL-UniRef50 (Prot-T5), that was pre-trained on BFD and finetuned on UniRef50 [66]. Both embeddings were computed using the `bio-embeddings` Python library [67]. Moreover, we used SeqVec [68] embeddings, obtained also using the `bio-embeddings` library as well as ESM-2 embeddings [69] obtained using the model file `esm2_t36_3B_UR50D` from ESM GitHub repository [70]. Both SeqVec and ESM-2 pLMs were pretrained on the UniRef50 dataset. For all the above pLMs, the encoder part of the model was used to compute the embeddings which were extracted from the last layer of the encoder. This represents the standard strategy used for evaluating the pre-trained embeddings on downstream tasks in the original papers [64, 68, 69]. Further information about embeddings, such as the number of parameters and embedding dimension, can be found in the supplementary table 5.

### 2.3 AA Index

PLMs are context-aware, resulting in different feature vectors for the same amino acid in different sequential contexts. To test the effect of information propagation through the protein graph (see section ***How much does the GNN architecture contribute to the performance?***), we also generated context-independent feature vectors, i.e., vectors whose values are not dependent on the neighborhood, serving as good baseline node features for our GNN models. For that purpose, we used the AAIndex database [15], a large collection of physicochemical and biochemical properties of amino acids. Using the AAIndex database, we constructed node features by collecting all returned properties of the respective amino acid into one vector. We used the Python AAIndex library [71] to extract AAIndex features. The AAIndex features were normalized over all amino acids, resulting in 566-dimensional feature vectors.

### 2.4 Datasets

As our main dataset, we used a benchmark designed by Yu et al. [10] involving 12 different ligands to build and test our models. Second, to validate that our methodology is on par with recent GNN-based approaches, we evaluated it on another dataset for protein-DNA and protein-RNA binding sites from the works of GraphBind [56] and GraphSite [57], details of which are given in Supplementary table 7.

The benchmarking dataset designed by Yu et al. [10] contains training and independent test sets of protein sequences and their corresponding actual binding residues for 12 different ligands, which include: 5 nucleotides (AMP, ADP, ATP, GTP, GDP), 5 ions (CA, MG, MN, FE, ZN), DNA, and HEME.

As the benchmark was used to test several sequence-based methods such as [10] and [33], and given that our method has a structural component, we needed to collect the corresponding 3D structures of the protein sequences. To achieve this, we downloaded the entire BioLip dataset [72], which was used to construct the benchmark, and we extracted the tertiary structures of the sequences by matching their PDB IDs and chain IDs. For sequences whose corresponding structures were not found in BioLip, we used the latest version of PDB [73] to extract the structures.

The PDB files were first parsed by the Biopython library [74] in order to obtain the sequences and the atomic coordinates. Some of the sequences obtained from the Biopython parser underwent minor manual corrections to match them with the sequences from the benchmark dataset. In total, the letters of some modified residues were changed for 12 sequences, one residue was skipped for 13 sequences, and 2 sequences were skipped due to a high mismatch between the sequence retrieved from the benchmark and the sequence retrieved after processing the PDB file. Finally, each residue from a sequence was associated with a 3D coordinate. The obtained coordinates were used to construct the protein graphs as described in section ***Protein graph construction*** using the Python Deep Graph Library (DGL) [75].

We also need to note that due to technical problems with the ProtT5 embeddings, we could not obtain embeddings for all of the proteins. In total, we could not obtain the protein graphs for 31 sequences. The sequences for which we could not generate the embeddings consisted of training sequences only, so this issue did not affect the reported results, as those were based on the test sets. Table 1 illustrates statistics of the benchmark datasets as well as the number of obtained protein graphs after the preprocessing phase.

**Table 1:** Yu benchmark summary.

| Ligand | Training sets |  |  |  | Independent test sets |  |  |
| --- | --- | --- | --- | --- | --- | --- | --- |
|  | Sequences | Missing protein graphs | Binding residues | Non-Binding residues | Sequences | Binding residues | Non-Binding residues |
| ATP | 221 | 0 | 3021 | 72334 | 50 | 647 | 16639 |
| ADP | 296 | 0 | 3833 | 98740 | 47 | 686 | 20327 |
| AMP | 145 | 0 | 1603 | 44401 | 33 | 392 | 10355 |
| GDP | 82 | 0 | 1101 | 26244 | 14 | 194 | 4180 |
| GTP | 54 | 1 | 745 | 21205 | 7 | 89 | 1868 |
| Ca<sup>2+</sup> | 965 | 4 | 4914 | 287801 | 165 | 785 | 53779 |
| Zn<sup>2+</sup> | 1168 | 16 | 4705 | 315235 | 176 | 744 | 47851 |
| Mg<sup>2+</sup> | 1138 | 7 | 3860 | 350716 | 217 | 852 | 72002 |
| Mn<sup>2+</sup> | 335 | 1 | 1496 | 112312 | 58 | 237 | 17484 |
| Fe<sup>3+</sup> | 173 | 1 | 818 | 50453 | 26 | 120 | 9092 |
| DNA | 335 | 0 | 6461 | 71320 | 52 | 973 | 16225 |
| HEME | 206 | 1 | 4380 | 49768 | 27 | 580 | 8630 |

### 2.5 Model hyperparameters

For building our models, we used the implementation of GCN and GAT provided by the Python library DGL-LifeSci [76], and we’ve trained and evaluated the models using the Pytorch Python library [77]. Our GCN architecture consisted of graph convolutional layers of size 512 with ReLU activation, a dropout rate of 0.5 [78], residual connections [79] and batch normalization [80]. At the same time, our GAT architecture consisted of graph attention layers of size 512, ReLU activations, dropout rate of 0.5, 4 attention heads, and residual connections. We used a dense layer with two softmax units on top of the GCN and GAT models to compute the node-level outputs. We also utilized a weighted version of the binary cross-entropy loss due to the high class imbalance of the datasets, the AdamW optimizer [81] as the optimization algorithm with `learning_rate=3e-4` and `weight_decay=1e-5`, and we trained all the models for `2000 epochs` with a `batch_size=32`. Since the process of training and evaluating the GNN models on the pLM embeddings is time-consuming, the hyperparameters of the GNN models were chosen after manual tuning on a random validation split from the training set. The range of values tried in the manual tuning is described in Supplementary table 6.

Regarding the sequence baseline models, we compared three model classes : Multi-Layer Perceptron (MLP), Random Forest (RF) and Support Vector Machines (SVM). The models were built using the embeddings from the ProtT5 language model. To select the sequence baseline architecture that will be used in the remaining experiments, we have performed 5-fold Cross-Validation (CV) on the ADP ligand training set using different hyperparameters of the model classes. The results of the 5-fold CV can be found in supplementary table 4. The SVM and RF models were implemented using the Scikit-learn Python library [82]. Moreover, the MLP classifiers were trained using the Pytorch Python library for 2000 epochs with a batch size of 32, and the reported validation scores of the MLPs represent the best validation scores obtained during the 2000 epochs training. To account for class imbalance in the sequence baselines, we used weighted binary cross-entropy as the loss function for the MLPs, and we assigned the `class_weight` parameter to `’balanced’` in the Scikit-learn implementation of the RF and SVM. Based on the 5-fold CV results, we have chosen the sequence baseline model in all remaining experiments to be a single-layer MLP with 512 units and with a dropout rate of 0.1, as it has the best mean CV score.

## 3 Results and Discussion

To evaluate the residue-level predictions of our models, we used standard binary classification metrics. Specifically, we have chosen to show our results with respect to the Matthews Correlation Coefficient (MCC) due to the significant class imbalance present in the datasets, as it has been shown that the MCC metric is one of the most suitable metrics in such cases [83].

Our recent work [33] shows that more complex LMs often yield better performance. Therefore we used the ProtT5 embeddings in most of our experiments as one of the most complex pLMs.

We used a random split of the processed benchmark training sets to obtain training and validation sets. The training/validation split ratio was designed for the validation sets to have the same size as the independent test sets. The validation sets were used to define the early stopping epoch while training the models. The training was stopped at the epoch with the best validation MCC. In the subsequent sections, we report the results of the independent test sets.

### 3.1 Effect of the number of graph convolutional layers

The effect of information propagation through the protein graph can best be seen by varying the number of convolutional layers. One round of graph convolution collects information from the neighborhood of a given node. Thus, as the number of graph convolutions increases, a given node will have access to more distant neighbors since the one-hop neighbors will already contain information about farther neighbors in their hidden features computed from previous rounds of graph convolution. Therefore, increasing the number of convolutional layers enables information propagation between distant parts of the graph. To test the effect of the number of convolutional layers on the prediction performance, we used graphs constructed using 6 ^°^A cutoff distance, ProtT5 embeddings, and we varied the number of graph convolutional layers in our standard GCN architecture; specifically we tried 1, 2, 4 and 6 layers. Furthermore, we report the mean and standard deviation of the MCC score for 5-fold cross-validation splits. The results are shown in figure 3 which was created using the supplementary table 1. The reported validation scores represent the best validation score obtained while training the models for 2000 epochs.

**Fig. 3:**
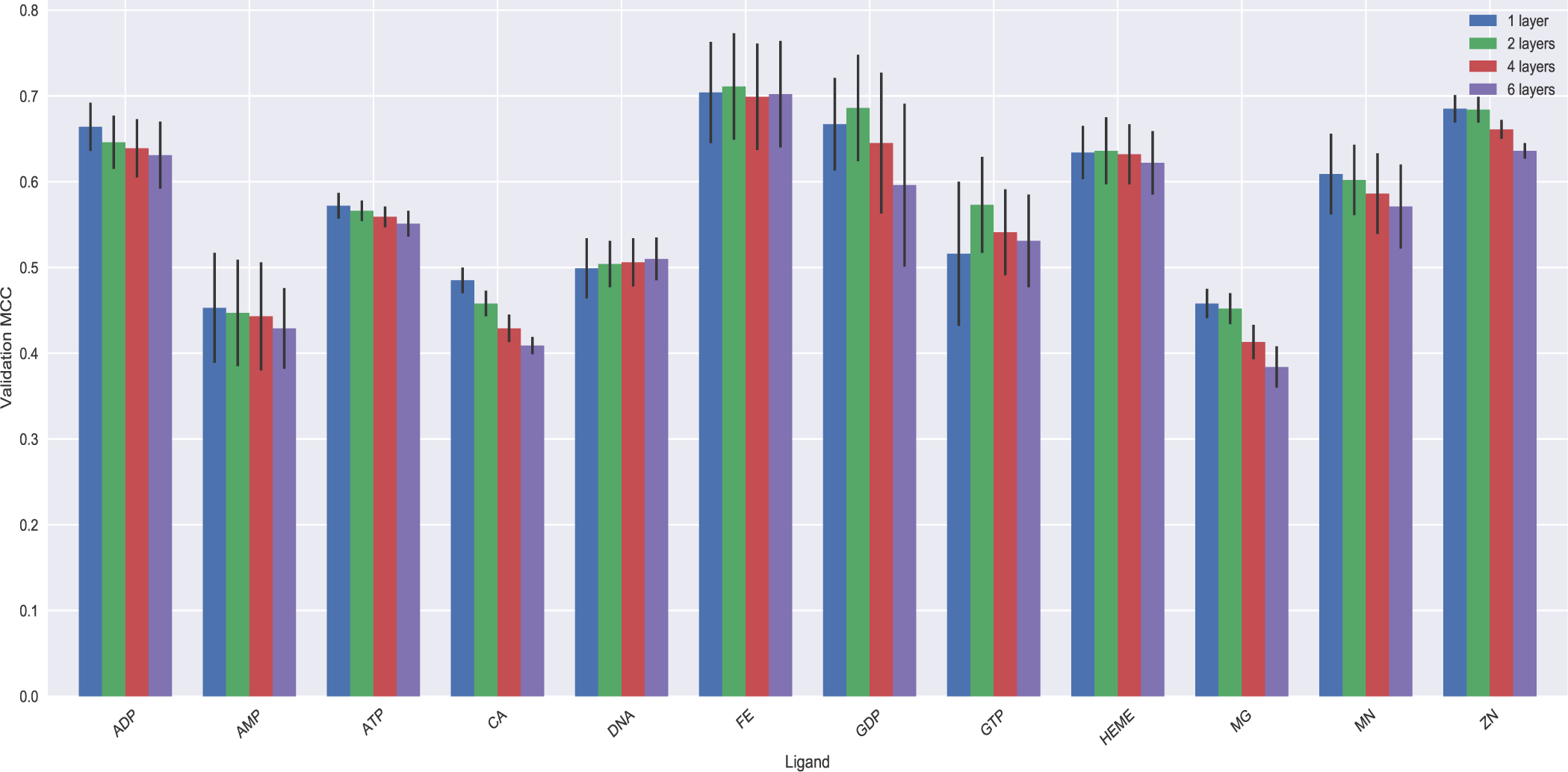
Effect of the number of graph convolutional layers. The bars represent the mean of the validation MCC scores for 5-fold Cross Validation splits. The error bars represent the standard deviation of the validation MCC scores. The colors correspond to the number of graph convolutional layers of 512 units.

We can observe that for about half of the ligand datasets the models constructed using different number of convolutional layers have very similar performance. Moreover, for most of the remaining ligand datasets, adding more graph convolutional layers decreases the performance. This suggests that there is little positive effect of adding more graph convolutional layers.

Based on the above observations, we decided to use a single-layered GNN architecture and an arbitrary random split with the same random seed in the remaining experiments. Another reason for choosing a single layer in the following experiments is to avoid the common oversmoothing problem in GNNs [84], where deep GNNs result in nearly indistinguishable node features in the last layers of the network, which may result in a poor performance in downstream tasks.

### 3.2 Effect of graph cutoff distances

Next, we tested the effect of the graph cutoff distance. The cutoff distance influences the graph’s connectivity as a higher cutoff distance results in more connections and thus leads to a more densely connected graph. In such a graph, a given node has more neighbors, and therefore more nodes are taken into account in information propagation to determine the state of the given node. A typical cutoff seen in other works is 6 ^°^A computed based on the distance of alpha carbons [85]. This work tested the following cutoff distances: 4 ^°^A, 6 ^°^A, 8 ^°^A, and 10 ^°^A. Moreover, we constructed an ensemble model using models trained on graphs built using the above cutoff distances. This model combines the predicted binary classes from each cutoff distance and outputs the most often observed class. An ensemble model that uses multiple cutoff distances removes the bias of choosing a predefined cutoff distance. Therefore it has the potential to improve the generalization capability of the GNN.

Table 2, compares the different cutoff distances and the ensemble model. We can observe that although the graph cutoff distance significantly affects the performance of the GCN model, there is no observable consistent trend by varying the cutoff distance. For instance, for some ligands such as ADP and HEME, a low cutoff distance (4 ^°^A) results in higher performance of the GCN, while for other ligands such as CA and MG, a high cutoff distance (10 ^°^A) improves the performance. Moreover, supplementary table 3 shows the effect of cutoff distances across multiple classification metrics, namely MCC, together with Precision and Recall.

**Table 2:** Effect of graph cutoff distance and the graph attention mechanism.

| Ligand | GCN |  |  |  |  | GAT |  |  |  |  | Sequence Baseline |
| --- | --- | --- | --- | --- | --- | --- | --- | --- | --- | --- | --- |
|  | 4 Å | 6 Å | 8 Å | 10 Å | Ensemble | 4 Å | 6 Å | 8 Å | 10 Å | Ensemble | Baseline |
| ADP | 0.569 | 0.564 | 0.581 | 0.557 | 0.584 | 0.571 | 0.578 | **0.597** | 0.582 | 0.583 | 0.553 |
| AMP | 0.450 | 0.412 | 0.424 | 0.419 | 0.445 | 0.449 | 0.463 | **0.489** | 0.475 | 0.482 | 0.416 |
| ATP | 0.546 | 0.537 | 0.538 | 0.557 | 0.569 | 0.566 | 0.575 | 0.572 | **0.587** | 0.583 | 0.501 |
| CA | 0.396 | 0.382 | 0.403 | 0.420 | 0.421 | 0.383 | 0.408 | 0.408 | 0.411 | 0.426 | **0.513** |
| DNA | 0.473 | 0.476 | 0.470 | 0.459 | 0.490 | 0.460 | 0.483 | **0.510** | 0.488 | 0.499 | 0.371 |
| FE | 0.618 | 0.645 | 0.614 | 0.645 | 0.645 | 0.704 | 0.668 | 0.692 | **0.719** | 0.703 | 0.651 |
| GDP | 0.665 | 0.668 | 0.737 | 0.693 | 0.710 | 0.696 | 0.695 | **0.746** | 0.705 | 0.744 | 0.651 |
| GTP | 0.537 | 0.514 | 0.575 | 0.564 | 0.556 | 0.666 | 0.669 | 0.670 | 0.573 | **0.695** | 0.524 |
| HEME | 0.689 | 0.672 | 0.736 | 0.675 | 0.691 | 0.675 | 0.674 | **0.743** | 0.682 | 0.685 | 0.720 |
| MG | 0.343 | 0.344 | 0.351 | 0.362 | **0.365** | 0.325 | 0.347 | 0.364 | 0.349 | 0.364 | 0.332 |
| MN | 0.617 | 0.606 | 0.594 | 0.590 | 0.634 | 0.602 | 0.642 | 0.607 | **0.642** | 0.638 | 0.585 |
| ZN | 0.660 | 0.681 | 0.673 | 0.693 | 0.699 | 0.670 | 0.672 | 0.685 | 0.690 | **0.699** | 0.671 |
| Average | 0.547 | 0.542 | 0.558 | 0.553 | **0.567** | 0.564 | 0.573 | 0.590 | 0.575 | **0.592** | 0.541 |

### 3.3 Effect of graph attention mechanism

For GAT, we tested the effect of the graph attention mechanism initially designed as a regularization strategy for the GNN models. The attention may contribute to a better generalization performance as the model attends only to relevant parts of the neighborhood of a node. To test the added value of the graph attention mechanism, we compared our shallow GCN model, with a shallow version of GAT where we used our standard GAT architecture with a single graph attention layer. Table 2 compares the GCN and GAT models for the different cutoff distances.

We see that, unlike in the case of GCN, for most datasets, there is a consistent improvement in the performance of the GAT model with increasing cutoff distance. This observation can be explained by the capacity of the attention mechanism to reduce noise in larger neighborhoods. For graphs obtained using a high cutoff distance, each node has a bigger neighborhood and collects information from more (distant) neighbors. Without using the attention mechanism, the model does not have the capacity to filter out irrelevant information. The graph attention mechanism fixes this issue by adjusting the neighbor weights to attend only to neighbors relevant for making the prediction.

Moreover, we observe that for all ligands and both for GAT and GCN, the ensemble models have better average performance across ligand datasets than all cutoff distances, and this performance is very similar to the average performance of the model with cutoff 8. These observations may suggest that the model with cutoff 8 can be considered as a lightweight proxy for the ensemble model in terms of the number of parameters and the required preprocessing steps. We will call those models GCN8 and GAT8 in the rest of the work. Table 2 shows that GAT8 has significantly higher performance than the GCN8 for the GTP ligand, while it is slightly more performant for most other ligands. Furthermore, the GAT8 significantly outperforms the sequence baseline for most ligands. In the following experiments, we, therefore, consider our best-performing model architecture to be the GAT8. Supplementary table 2 then also includes a comparison of GAT and GCN using more classification metrics.

### 3.4 What is the attention attentive to?

In the previous section, we showed that attention helps to improve the accuracy of predictions in comparison with GCN; we were further wondering what amino acids were helpful and therefore investigated a number of binding sites of Zn ion, GTP and HEME as three variable representatives of studied ligands. We were specifically investigating cases where a ligand-binding residue was not predicted by GCN but was correctly predicted with GAT. To do that, we used 10 ^°^A protein graphs; for every binding residue, we extracted the attention value for each neighbor. As our model uses four attention heads, the attention values were averaged across the heads. Then, individually for each binding residue, we colored the binding site with a relative contribution of attention of the binding residue neighbors. We observed that in many cases, the residues with the highest attention were the other ligand-binding residues (and sequence neighbors of the studied residue). In many cases, the binding residues were often physically close to the ligand, but we also observed cases where the residues with the highest attention were on the other part of the binding site and away from the studied residue (see Figure 4).

**Fig. 4:**
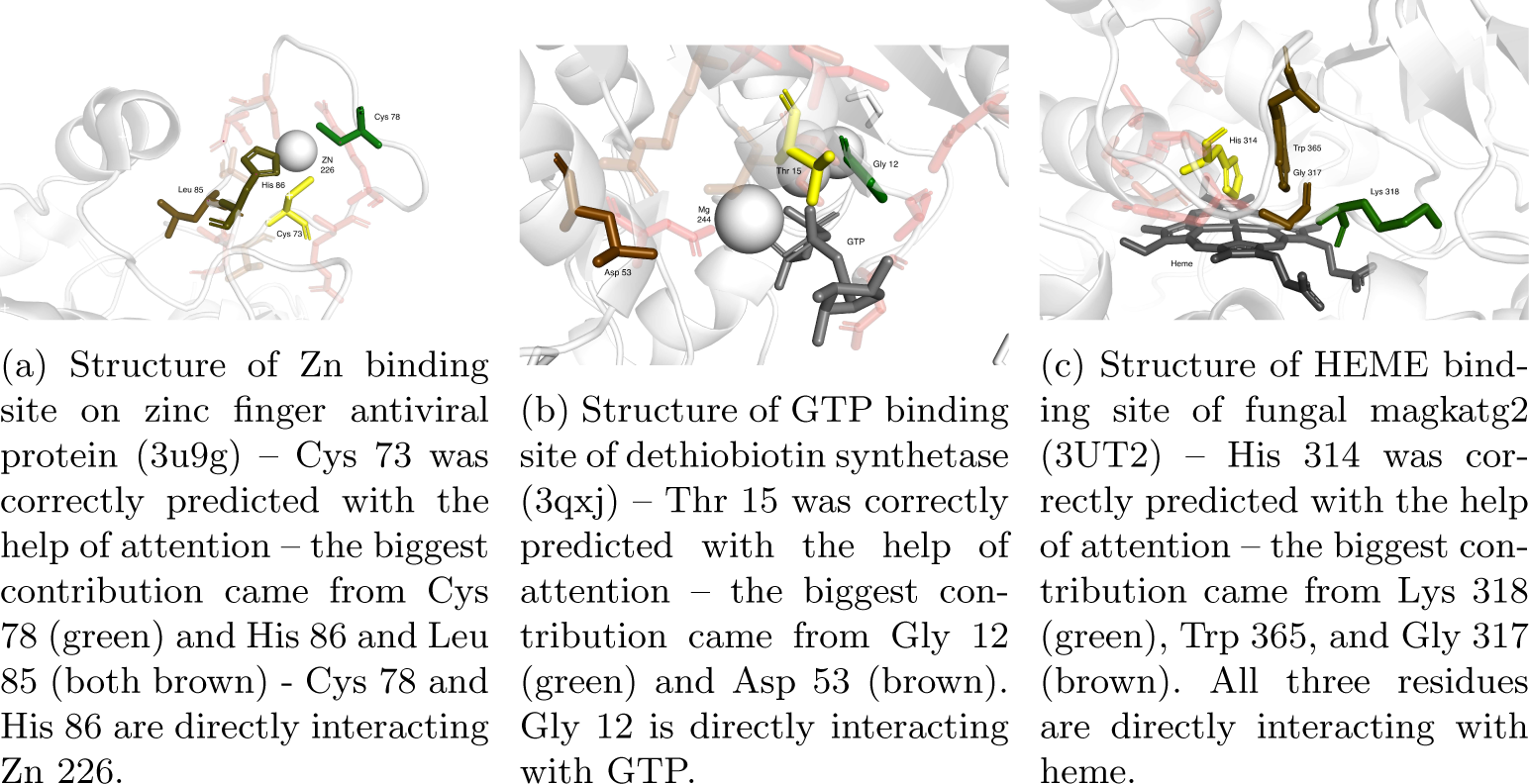
Visualization of the attention. The binding residue and its neighbors are represented as sticks. The binding residue is colored yellow with neighbors going from green (highest attention) to red (lowest attention). The low-attention neighbors are partially transparent. The ligand is colored gray.

We should emphasize that the goal of this exercise was to offer a visual way of inspecting the attention, but a more quantitative approach should be taken to draw a conclusive statement regarding the attention. This is further supported by the fact that we also encountered instances where it was not clear how could the residues with high attention contributed to the accurate prediction of the studied residue.

### 3.5 Comparison with existing methods

To put the proposed approach in the context of existing research, we compared our GAT8 model with the Prot-T5 embeddings, which consistently demonstrated higher performance in the previous experiments, to other approaches which were trained and tested using the Yu benchmark dataset. Namely, TargetS [10], EC-RUS [86], and SXG-Bsite [87], which are based on different hand-crafted, but context-dependent features as described in section 1. For each of the three methods, we show the results of the best-performing versions of those methods as presented in the respective papers. Table 3 compares the methods using the area under ROC curve (ROC-AUC) and MCC. Our GAT8 model with ProtT5 embeddings outperforms all of the methods on the MCC metric for all ligand datasets, and on the ROC-AUC metric for most datasets. However, it should be emphasized that the presented methods are sequence-based, using only predicted structural features (such as predicted secondary structure). On the other hand, the presented approach does not incorporate 3D structure directly, as the protein graph only approximates the 3D information.

**Table 3:** Comparison with existing methods - Yu benchmark.

| Ligand | AUC |  |  |  | MCC |  |  |  |
| --- | --- | --- | --- | --- | --- | --- | --- | --- |
|  | TargetS | EC-RUS | SXGBsite | ProtT5 GAT8 | TargetS | EC-RUS | SXGBsite | ProtT5 GAT8 |
| ADP | 0.896 | 0.872 | 0.907 | **0.945** | 0.507 | 0.511 | 0.521 | **0.597** |
| AMP | 0.83 | 0.815 | 0.851 | **0.892** | 0.359 | 0.393 | 0.366 | **0.489** |
| ATP | 0.898 | 0.871 | 0.886 | **0.936** | 0.502 | 0.506 | 0.448 | **0.572** |
| CA | 0.767 | 0.77 | 0.757 | **0.882** | 0.243 | 0.225 | 0.167 | **0.408** |
| DNA | 0.836 | 0.814 | 0.827 | **0.932** | 0.377 | 0.319 | 0.27 | **0.510** |
| FE | 0.945 | 0.936 | 0.913 | **0.986** | 0.479 | 0.49 | 0.454 | **0.692** |
| GDP | 0.896 | 0.872 | 0.93 | **0.963** | 0.55 | 0.579 | 0.678 | **0.746** |
| GTP | 0.855 | 0.861 | 0.883 | **0.932** | 0.617 | 0.641 | 0.572 | **0.670** |
| HEME | 0.907 | 0.935 | 0.9 | **0.976** | 0.598 | 0.64 | 0.555 | **0.743** |
| MG | 0.706 | 0.78 | **0.819** | 0.782 | 0.294 | 0.317 | 0.326 | **0.364** |
| MN | 0.888 | 0.891 | 0.888 | **0.920** | 0.449 | 0.31 | 0.329 | **0.607** |
| ZN | 0.936 | 0.958 | 0.892 | **0.962** | 0.527 | 0.437 | 0.363 | **0.685** |

Finally, we also validate that our approach is comparable with recently published methods that predict nucleic acid binding using GNNs. Specifically, supplementary table 8 compares our approach with GraphBind [56] and GraphSite [57] which used variations of GNNs, in addition to GeoBind [88] and EquiPNAS [89] that used combinations of GNNs and PLMs.For each of the presented methods, we report the best-performing version which relied on the experimental protein structure to construct the protein graph. We compare the methods using the area under ROC curve (AUC), the area under the Precision-Recall curve (AUPR), and MCC. We report the scores directly from the original published results of the methods whenever the score was available. While EquiPNAS and GeoBind showed the best performance on the DNA/RNA benchmarks, our GAT8 model with ProtT5 embeddings shows similar performance to GraphBind and GraphSite especially on the DNA benchmark.

### 3.6 How much does the GNN architecture contribute to the performance?

In the previous sections, we observed that the GNN architecture improves the performance of the ProtT5 pLM. This observation prompted us to the necessity of quantifying how much the structural information processed by the GNN architecture contributes to the predictive performance of sequence-based pLMs. To this end, we designed two experiments to analyze the interplay of sequence information represented by node embeddings and structural information embedded in the graph connectivity. The first experiment involved comparing the GAT8 architecture with the sequence baseline model for several node embeddings using the Yu benchmark. Specifically, we compared one embedding with context-independent features and four embeddings from four different pLMs. The first embedding uses the context-independent AAIndex physico-chemical properties of amino acids, where a residue is represented by the same feature vector independently of its sequential context. The four remaining models use different context-aware pLMs of varying complexity. SeqVec embeddings and Prot-BERT, which are relatively less complex, as well as ProtT5 and ESM-2 embeddings, which are relatively more complex. The embedding complexity can be measured by two main indicators: the number of parameters of the pLM, and the dimensionality of the embedding (see supplementary table 5). The model complexity increases when one or both indicators increase. We measured the effect of the structure information by calculating the absolute (absolute Δ) and relative improvements (relative Δ) of the GAT8 models over their respective sequence baselines. The different test MCC scores and their respective absolute and relative improvements are presented in table 4. A more detailed comparison using more classification metrics between the embeddings is available in supplementary table 3.

**Table 4:** Effect of different embeddings. The relative improvement over the sequence baseline in the MCC score is computed as the GAT8 model’s MCC score minus the sequence baseline’s MCC score, divided by the MCC score of the sequence baseline.

| Embedding | Ligand | ADP | AMP | ATP | CA | DNA | FE | GDP | GTP | HEME | MG | MN | ZN | Average |
| --- | --- | --- | --- | --- | --- | --- | --- | --- | --- | --- | --- | --- | --- | --- |
| **AAIndex** | Sequence | 0.067 | 0.057 | 0.065 | 0.115 | 0.122 | 0.180 | 0.106 | 0.108 | 0.116 | 0.065 | 0.170 | 0.257 | 0.119 |
|  | GAT8 | 0.142 | 0.101 | 0.106 | 0.122 | 0.168 | 0.241 | 0.215 | 0.192 | 0.214 | 0.083 | 0.204 | 0.331 | 0.177 |
| | Absolute $\Delta$ | 0.075 | 0.044 | 0.041 | 0.007 | 0.046 | 0.061 | 0.109 | 0.084 | 0.098 | 0.018 | 0.034 | 0.074 | 0.058 |
| | Relative $\Delta$ | 1.119 | 0.772 | 0.631 | 0.061 | 0.377 | 0.339 | 1.028 | 0.778 | 0.845 | 0.277 | 0.200 | 0.288 | 0.560 |
| **SeqVec** | Sequence | 0.519 | 0.266 | 0.428 | 0.286 | 0.254 | 0.498 | 0.580 | 0.498 | 0.523 | 0.243 | 0.423 | 0.537 | 0.421 |
|  | GAT8 | 0.571 | 0.365 | 0.512 | 0.310 | 0.322 | 0.587 | 0.640 | 0.613 | 0.602 | 0.298 | 0.480 | 0.598 | 0.492 |
| | Absolute $\Delta$ | 0.052 | 0.099 | 0.084 | 0.024 | 0.068 | 0.089 | 0.060 | 0.115 | 0.079 | 0.055 | 0.057 | 0.061 | 0.070 |
| | Relative $\Delta$ | 0.101 | 0.370 | 0.196 | 0.084 | 0.266 | 0.179 | 0.104 | 0.230 | 0.150 | 0.226 | 0.135 | 0.115 | 0.180 |
| **ProtBERT** | Sequence | 0.426 | 0.286 | 0.381 | 0.298 | 0.348 | 0.594 | 0.543 | 0.405 | 0.477 | 0.287 | 0.437 | 0.573 | 0.421 |
|  | GAT8 | 0.504 | 0.326 | 0.445 | 0.378 | 0.381 | 0.635 | 0.580 | 0.552 | 0.564 | 0.324 | 0.514 | 0.618 | 0.485 |
| | Absolute $\Delta$ | 0.078 | 0.040 | 0.064 | 0.080 | 0.033 | 0.041 | 0.037 | 0.147 | 0.087 | 0.037 | 0.077 | 0.045 | 0.064 |
| | Relative $\Delta$ | 0.184 | 0.140 | 0.167 | 0.268 | 0.096 | 0.069 | 0.067 | 0.362 | 0.183 | 0.128 | 0.175 | 0.078 | 0.160 |
| **ProtT5** | Sequence | 0.553 | 0.416 | 0.501 | 0.513 | 0.371 | 0.651 | 0.651 | 0.524 | 0.720 | 0.332 | 0.585 | 0.671 | 0.541 |
|  | GAT8 | 0.597 | 0.489 | 0.572 | 0.408 | 0.510 | 0.692 | 0.746 | 0.670 | 0.743 | 0.364 | 0.607 | 0.649 | 0.587 |
| | Absolute $\Delta$ | 0.044 | 0.073 | 0.071 | -0.105 | 0.139 | 0.041 | 0.095 | 0.146 | 0.023 | 0.032 | 0.022 | -0.023 | 0.047 |
| | Relative $\Delta$ | 0.080 | 0.176 | 0.142 | -0.205 | 0.376 | 0.063 | 0.146 | 0.277 | 0.032 | 0.097 | 0.038 | -0.034 | 0.099 |
| **ESM-2** | Sequence | 0.570 | 0.476 | 0.540 | 0.382 | 0.462 | 0.641 | 0.702 | 0.677 | 0.722 | 0.309 | 0.576 | 0.647 | 0.559 |
|  | GAT8 | 0.616 | 0.493 | 0.597 | 0.401 | 0.647 | 0.643 | 0.750 | 0.671 | 0.755 | 0.350 | 0.597 | 0.683 | 0.600 |
| | Absolute $\Delta$ | 0.046 | 0.017 | 0.057 | 0.019 | 0.185 | 0.002 | 0.048 | -0.006 | 0.033 | 0.041 | 0.021 | 0.036 | 0.042 |
| | Relative $\Delta$ | 0.082 | 0.036 | 0.106 | 0.051 | 0.401 | 0.002 | 0.069 | -0.009 | 0.046 | 0.132 | 0.036 | 0.056 | 0.084 |

Table 4 indicates that the absolute improvement in the test MCC score of the GAT8 model over the sequence baseline is positive on average across all ligand datasets. Moreover, although different embeddings have varying degrees of absolute improvement depending on the ligand dataset, they have similar values on average. Nevertheless, in the case of the less complex AAIndex, SeqVec and ProtBERT embeddings, we can observe the lowest relative improvements in the MCC score of the GAT8 model over the sequence baseline for most ligands, while the more complex ESM-2 and ProtT5 embeddings show smaller relative improvements. These observations show that while the protein structural information almost always improves the sequence baseline regardless of the chosen embedding, the relative effect is more pronounced for simple embeddings and decreases with the complexity of the language models.

To consolidate the relative improvement observations, we have performed statistical significance tests in a 5-fold cross-validation setting for all embeddings and across all ligand datasets. Table 5 presents the results of t-tests for the mean relative improvement scores of the GAT8 model over the sequence baseline in a 5-fold CV setting. The null hypothesis of the t-tests is that there is no relative improvement, and the significance threshold is chosen to be 0.01. Table 5, shows that for all embeddings, most relative improvement values are statistically significant (*P − value <* 0.01.)

**Table 5:** Statistical significance tests for relative improvement values. The mean and standard deviation of the relative improvements are computed from GAT8 MCC scores and sequence baseline MCC scores of the validation sets from 5-fold CV. The P-values correspond to the result of the t-test performed on the relative improvement values from the CV folds. Statistically significant P-values are displayed in bold (*P −value <* 0.01)

| Embedding | Ligand | ADP | AMP | ATP | CA | DNA | FE | GDP | GTP | HEME | MG | MN | ZN |
| --- | --- | --- | --- | --- | --- | --- | --- | --- | --- | --- | --- | --- | --- |
| AAIndex | Relative $\Delta$ | 1.266<br>$\pm 0.336$ | 0.451<br>$\pm 0.211$ | 0.851<br>$\pm 0.141$ | 0.224<br>$\pm 0.071$ | 0.367<br>$\pm 0.05$ | 0.423<br>$\pm 0.111$ | 1.144<br>$\pm 0.134$ | 0.815<br>$\pm 0.382$ | 0.572<br>$\pm 0.225$ | 0.252<br>$\pm 0.05$ | 0.198<br>$\pm 0.105$ | 0.284<br>$\pm 0.012$ |
|  | P-Value | **0.0011** | **0.0087** | **0.0002** | **0.0021** | **0.0001** | **0.0010** | **0.0000** | **0.0088** | **0.0048** | **0.0003** | 0.0135 | **0.0000** |
| SeqVec | Relative $\Delta$ | 0.111<br>$\pm 0.034$ | 0.309<br>$\pm 0.091$ | 0.18<br>$\pm 0.076$ | 0.119<br>$\pm 0.047$ | 0.235<br>$\pm 0.063$ | 0.28<br>$\pm 0.037$ | 0.158<br>$\pm 0.07$ | 0.303<br>$\pm 0.09$ | 0.122<br>$\pm 0.031$ | 0.119<br>$\pm 0.027$ | 0.247<br>$\pm 0.066$ | 0.107<br>$\pm 0.024$ |
|  | P-Value | **0.0019** | **0.0016** | **0.0062** | **0.0048** | **0.0011** | **0.0001** | **0.0072** | **0.0017** | **0.0010** | **0.0006** | **0.0011** | **0.0006** |
| ProtBERT | Relative $\Delta$ | 0.201<br>$\pm 0.034$ | 0.18<br>$\pm 0.058$ | 0.222<br>$\pm 0.09$ | 0.188<br>$\pm 0.028$ | 0.128<br>$\pm 0.027$ | 0.079<br>$\pm 0.124$ | 0.186<br>$\pm 0.042$ | 0.328<br>$\pm 0.16$ | 0.148<br>$\pm 0.015$ | 0.164<br>$\pm 0.053$ | 0.138<br>$\pm 0.036$ | 0.108<br>$\pm 0.014$ |
|  | P-Value | **0.0002** | **0.0023** | **0.0053** | **0.0001** | **0.0005** | 0.2278 | **0.0006** | 0.0102 | **0.0000** | **0.0023** | **0.0010** | **0.0001** |
| ProtT5 | Relative $\Delta$ | 0.069<br>$\pm 0.017$ | 0.127<br>$\pm 0.127$ | 0.109<br>$\pm 0.027$ | 0.103<br>$\pm 0.037$ | 0.066<br>$\pm 0.034$ | 0.065<br>$\pm 0.114$ | 0.107<br>$\pm 0.052$ | 0.218<br>$\pm 0.084$ | 0.057<br>$\pm 0.019$ | 0.059<br>$\pm 0.018$ | 0.064<br>$\pm 0.027$ | 0.032<br>$\pm 0.012$ |
|  | P-Value | **0.0008** | 0.0902 | **0.0009** | **0.0034** | 0.0118 | 0.2674 | 0.0101 | **0.0043** | **0.0025** | **0.0018** | **0.0062** | **0.0044** |
| ESM-2 | Relative $\Delta$ | 0.069<br>$\pm 0.008$ | 0.126<br>$\pm 0.037$ | 0.07<br>$\pm 0.012$ | 0.101<br>$\pm 0.033$ | 0.035<br>$\pm 0.028$ | 0.026<br>$\pm 0.013$ | 0.033<br>$\pm 0.014$ | 0.056<br>$\pm 0.051$ | 0.031<br>$\pm 0.019$ | 0.097<br>$\pm 0.032$ | 0.05<br>$\pm 0.023$ | 0.06<br>$\pm 0.014$ |
|  | P-Value | **0.0000** | **0.0015** | **0.0002** | **0.0024** | 0.0468 | 0.0117 | **0.0064** | 0.0676 | 0.0208 | **0.0024** | **0.0078** | **0.0007** |

To quantify how much improvement is caused by the concrete graph topology as opposed to random propagation of information, we devised the following experiment. We compared the GAT8 model with graphs constructed using the experimental PDB structure called “original” with a “random” version of the same model, where the original graph was replaced by a random graph with perturbed edges. Specifically, we randomly assigned edges between residues and explicitly removed every edge in the original graph. The “random” model provides a solid baseline against which to measure the effect of the experimental structure information in the GNN architecture and its relationship with pLMs. In table 6, we report the absolute and relative improvements in test MCC scores of the GAT8 model with original graphs over their respective random graph baselines. We observe from the absolute improvement scores that for all embeddings, the original structure almost always contributes positively to the performance. Nevertheless, this effect tends to decrease on average both in terms of absolute and relative improvement, especially for more complex pLMs.

**Table 6:** Effect of original structure. The relative improvement over the sequence baseline in the MCC score is computed as the GAT8 model’s MCC score minus the sequence baseline’s MCC score, divided by the MCC score of the sequence baseline.

| Embedding | Ligand | ADP | AMP | ATP | CA | DNA | FE | GDP | GTP | HEME | MG | MN | ZN | Average |
| --- | --- | --- | --- | --- | --- | --- | --- | --- | --- | --- | --- | --- | --- | --- |
| **AAIndex** | Original | 0.142 | 0.101 | 0.106 | 0.122 | 0.168 | 0.241 | 0.215 | 0.192 | 0.214 | 0.083 | 0.204 | 0.331 | 0.177 |
|  | Random | 0.052 | 0.054 | 0.064 | 0.115 | 0.122 | 0.193 | 0.083 | 0.090 | 0.135 | 0.066 | 0.165 | 0.256 | 0.116 |
| | Absolute $\Delta$ | 0.090 | 0.047 | 0.042 | 0.007 | 0.046 | 0.048 | 0.132 | 0.102 | 0.079 | 0.017 | 0.039 | 0.075 | 0.060 |
| | Relative $\Delta$ | 1.734 | 0.870 | 0.646 | 0.065 | 0.377 | 0.249 | 1.579 | 1.125 | 0.585 | 0.261 | 0.239 | 0.295 | 0.669 |
| **SeqVec** | Original | 0.571 | 0.365 | 0.512 | 0.310 | 0.322 | 0.587 | 0.640 | 0.613 | 0.602 | 0.298 | 0.480 | 0.598 | 0.492 |
|  | Random | 0.528 | 0.352 | 0.500 | 0.291 | 0.298 | 0.576 | 0.611 | 0.613 | 0.566 | 0.293 | 0.479 | 0.572 | 0.473 |
| | Absolute $\Delta$ | 0.043 | 0.013 | 0.012 | 0.019 | 0.024 | 0.011 | 0.029 | 0.000 | 0.036 | 0.005 | 0.001 | 0.026 | 0.018 |
| | Relative $\Delta$ | 0.082 | 0.036 | 0.024 | 0.064 | 0.082 | 0.018 | 0.048 | 0.000 | 0.064 | 0.017 | 0.002 | 0.045 | 0.040 |
| **ProtBERT** | Original | 0.504 | 0.326 | 0.445 | 0.378 | 0.381 | 0.635 | 0.580 | 0.552 | 0.564 | 0.324 | 0.514 | 0.618 | 0.485 |
|  | Random | 0.468 | 0.305 | 0.423 | 0.337 | 0.357 | 0.601 | 0.589 | 0.535 | 0.527 | 0.297 | 0.485 | 0.614 | 0.462 |
| | Absolute $\Delta$ | 0.036 | 0.021 | 0.022 | 0.041 | 0.024 | 0.034 | -0.009 | 0.017 | 0.037 | 0.027 | 0.029 | 0.004 | 0.024 |
| | Relative $\Delta$ | 0.078 | 0.068 | 0.051 | 0.121 | 0.067 | 0.057 | -0.016 | 0.031 | 0.070 | 0.090 | 0.060 | 0.007 | 0.057 |
| **ProtT5** | Original | 0.597 | 0.489 | 0.572 | 0.408 | 0.510 | 0.692 | 0.746 | 0.670 | 0.743 | 0.364 | 0.607 | 0.649 | 0.587 |
|  | Random | 0.583 | 0.471 | 0.560 | 0.381 | 0.494 | 0.688 | 0.706 | 0.635 | 0.729 | 0.351 | 0.589 | 0.689 | 0.573 |
| | Absolute $\Delta$ | 0.014 | 0.018 | 0.012 | 0.027 | 0.016 | 0.004 | 0.040 | 0.035 | 0.014 | 0.013 | 0.018 | -0.040 | 0.014 |
| | Relative $\Delta$ | 0.024 | 0.038 | 0.021 | 0.071 | 0.031 | 0.005 | 0.057 | 0.055 | 0.020 | 0.037 | 0.030 | -0.059 | 0.028 |
| **ESM-2** | Original | 0.616 | 0.493 | 0.597 | 0.401 | 0.647 | 0.643 | 0.750 | 0.671 | 0.755 | 0.350 | 0.597 | 0.683 | 0.600 |
|  | Random | 0.617 | 0.507 | 0.603 | 0.416 | 0.475 | 0.674 | 0.755 | 0.705 | 0.750 | 0.342 | 0.574 | 0.680 | 0.591 |
| | Absolute $\Delta$ | -0.001 | -0.014 | -0.006 | -0.015 | 0.172 | -0.031 | -0.005 | -0.034 | 0.005 | 0.008 | 0.023 | 0.003 | 0.009 |
| | Relative $\Delta$ | -0.002 | -0.027 | -0.010 | -0.035 | 0.363 | -0.046 | -0.007 | -0.048 | 0.006 | 0.023 | 0.040 | 0.005 | 0.022 |

The results of both experiments suggest that due to the fact that more complex embeddings significantly improve the performance of the sequence and the random graph baselines, a significant part of the structure information necessary for predicting protein-ligand binding sites is already encoded in the protein language models. This may be explained by the fact that as complex protein language models were built using masked language modeling, large number of parameters and huge training sets, important relationships between residues that correlate with structural features may already be captured in the embeddings and can thus be used for binding site predictions.

## 4 Conclusion

In this work, we integrated sequence-based and structure-based paradigms for predicting protein-ligand binding sites by designing a GNN model augmented with protein language model embeddings. While the model’s performance varies with the cutoff distance used to construct the protein graph, the introduction of the graph attention mechanism significantly enhances predictive performance for densely connected graphs. Our findings indicate that although the structural information processed by the GNN architecture generally contributes positively to the model’s performance, this effect is more pronounced with simple node features and diminishes with the use of more complex language models. Overall, our research demonstrates the potential utility of combining sequence-based and structure-based approaches—specifically, using a GNN model enhanced with protein language model embeddings—to improve protein-ligand binding site prediction. This is particularly promising given the increasing availability of predicted 3D models. Although slight inaccuracies in atom positions within these predicted structures might pose challenges for tasks like molecular docking, they should not significantly impact the protein-ligand residue prediction task. This is because the graph topology, which serves as the input to the GNN, is merely an approximation of the protein’s three-dimensional structure and remains relatively unaffected by minor perturbations in atom positions. Consequently, we believe that integrating protein sequence information from language models with 3D structure data is a promising approach for predicting protein-ligand binding residues.

## Declarations

### • Funding

This work was supported by the Czech Science Foundation grant 23-07349S. Computational resources were provided by the e-INFRA CZ project (ID:90254), supported by the Ministry of Education, Youth and Sports of the Czech Republic.

Part of this work was carried out with the support of ELIXIR CZ Research Infrastructure (ID LM2023055, MEYS CR).

### • Ethics approval

Not applicable.

### • Consent to participate

Not applicable.

### • Consent for publication

Not applicable.

### • Availability of data and materials

The datasets generated and/or analyzed during the current study are available in the following GitHub repository https://github.com/hamzagamouh/pt-lm-gnn.

### • Code availability

The source code that was used to generate the results of the current study is available in the following GitHub repository https://github.com/hamzagamouh/pt-lm-gnn.

### • Authors’ contributions

D.H. and H.G. conceived the project, D.H. supervised the project, H.G. implemented the method and conducted the experiments, H.G., D.H., and M.N. analyzed the results. H. G. wrote most of the manuscript, H.G., D.H., and M.N. reviewed the manuscript.

## 5 Supplementary material

**Supplementary Table 1:** Effect of the number of graph convolutional layers with ProtT5 embeddings and cutoff distance of 6 ^°^A. The displayed scores are means and standard deviations of validation MCC scores from 5-fold cross-validation.

| Ligand | Number of convolutional layers |  |  |  |
| --- | --- | --- | --- | --- |
|  | 1 | 2 | 4 | 6 |
| ADP | **0.664 ± 0.028** | 0.646 ± 0.031 | 0.639 ± 0.034 | 0.631 ± 0.039 |
| AMP | **0.453 ± 0.064** | 0.447 ± 0.062 | 0.443 ± 0.063 | 0.429 ± 0.047 |
| ATP | **0.572 ± 0.015** | 0.566 ± 0.012 | 0.559 ± 0.012 | 0.551 ± 0.015 |
| CA | **0.485 ± 0.015** | 0.458 ± 0.015 | 0.429 ± 0.016 | 0.409 ± 0.010 |
| DNA | 0.499 ± 0.035 | **0.504 ± 0.027** | 0.506 ± 0.028 | 0.510 ± 0.025 |
| FE | 0.704 ± 0.059 | **0.711 ± 0.062** | 0.699 ± 0.062 | 0.702 ± 0.062 |
| GDP | 0.667 ± 0.054 | **0.686 ± 0.062** | 0.645 ± 0.082 | 0.596 ± 0.095 |
| GTP | 0.516 ± 0.084 | **0.573 ± 0.056** | 0.541 ± 0.050 | 0.531 ± 0.054 |
| HEME | 0.634 ± 0.031 | **0.636 ± 0.039** | 0.632 ± 0.035 | 0.622 ± 0.037 |
| MG | **0.458 ± 0.017** | 0.452 ± 0.018 | 0.413 ± 0.020 | 0.384 ± 0.024 |
| MN | **0.609 ± 0.047** | 0.602 ± 0.041 | 0.586 ± 0.047 | 0.571 ± 0.049 |
| ZN | **0.685 ± 0.016** | 0.684 ± 0.015 | 0.661 ± 0.011 | 0.636 ± 0.009 |

**Supplementary Table 2:** Comparison of GAT and GCN for ProtT5 embeddings and for different cutoff distances.

| Ligand | Cutoff | GCN |  |  | GAT |  |  |
| --- | --- | --- | --- | --- | --- | --- | --- |
|  |  | Precision | Recall | MCC | Precision | Recall | MCC |
| ADP | 4 | 0.663 | 0.51 | 0.569 | 0.674 | 0.504 | 0.571 |
|  | 6 | 0.658 | 0.504 | 0.564 | 0.704 | 0.493 | 0.578 |
|  | 8 | 0.656 | 0.536 | 0.581 | 0.674 | 0.55 | 0.597 |
|  | 10 | 0.661 | 0.491 | 0.557 | 0.641 | 0.551 | 0.582 |
|  | Ensemble | 0.726 | 0.487 | 0.584 | 0.713 | 0.496 | 0.583 |
| AMP | 4 | 0.49 | 0.449 | 0.45 | 0.398 | 0.561 | 0.449 |
|  | 6 | 0.443 | 0.423 | 0.412 | 0.465 | 0.503 | 0.463 |
|  | 8 | 0.448 | 0.441 | 0.424 | 0.67 | 0.378 | 0.489 |
|  | 10 | 0.458 | 0.421 | 0.419 | 0.618 | 0.388 | 0.475 |
|  | Ensemble | 0.54 | 0.395 | 0.445 | 0.632 | 0.39 | 0.482 |
| ATP | 4 | 0.578 | 0.547 | 0.546 | 0.567 | 0.598 | 0.566 |
|  | 6 | 0.543 | 0.567 | 0.537 | 0.651 | 0.533 | 0.575 |
|  | 8 | 0.554 | 0.558 | 0.538 | 0.62 | 0.556 | 0.572 |
|  | 10 | 0.579 | 0.567 | 0.557 | 0.66 | 0.547 | 0.587 |
|  | Ensemble | 0.651 | 0.522 | 0.569 | 0.677 | 0.526 | 0.583 |
| CA | 4 | 0.526 | 0.308 | 0.396 | 0.473 | 0.322 | 0.383 |
|  | 6 | 0.531 | 0.284 | 0.382 | 0.522 | 0.329 | 0.408 |
|  | 8 | 0.551 | 0.303 | 0.403 | 0.533 | 0.322 | 0.408 |
|  | 10 | 0.595 | 0.304 | 0.42 | 0.524 | 0.332 | 0.411 |
|  | Ensemble | 0.647 | 0.28 | 0.421 | 0.613 | 0.303 | 0.426 |
| DNA | 4 | 0.48 | 0.53 | 0.473 | 0.442 | 0.552 | 0.46 |
|  | 6 | 0.476 | 0.541 | 0.476 | 0.459 | 0.579 | 0.483 |
|  | 8 | 0.411 | 0.625 | 0.47 | 0.438 | 0.674 | 0.51 |
|  | 10 | 0.43 | 0.568 | 0.459 | 0.474 | 0.569 | 0.488 |
|  | Ensemble | 0.519 | 0.518 | 0.49 | 0.5 | 0.559 | 0.499 |
| FE | 4 | 0.463 | 0.842 | 0.618 | 0.602 | 0.833 | 0.704 |
|  | 6 | 0.488 | 0.867 | 0.645 | 0.572 | 0.792 | 0.668 |
|  | 8 | 0.467 | 0.825 | 0.614 | 0.594 | 0.817 | 0.692 |
|  | 10 | 0.493 | 0.858 | 0.645 | 0.609 | 0.858 | 0.719 |
|  | Ensemble | 0.513 | 0.825 | 0.645 | 0.613 | 0.817 | 0.703 |
| GDP | 4 | 0.756 | 0.608 | 0.665 | 0.851 | 0.588 | 0.696 |
|  | 6 | 0.782 | 0.593 | 0.668 | 0.801 | 0.624 | 0.695 |
|  | 8 | 0.879 | 0.634 | 0.737 | 0.937 | 0.608 | 0.746 |
|  | 10 | 0.796 | 0.624 | 0.693 | 0.735 | 0.701 | 0.705 |
|  | Ensemble | 0.896 | 0.577 | 0.71 | 0.922 | 0.613 | 0.744 |
| GTP | 4 | 0.469 | 0.674 | 0.537 | 0.761 | 0.607 | 0.666 |
|  | 6 | 0.47 | 0.618 | 0.514 | 0.753 | 0.618 | 0.669 |
|  | 8 | 0.553 | 0.64 | 0.575 | 0.731 | 0.64 | 0.67 |
|  | 10 | 0.544 | 0.629 | 0.564 | 0.508 | 0.697 | 0.573 |
|  | Ensemble | 0.58 | 0.573 | 0.556 | 0.809 | 0.618 | 0.695 |
| HEME | 4 | 0.716 | 0.7 | 0.689 | 0.715 | 0.676 | 0.675 |
|  | 6 | 0.718 | 0.667 | 0.672 | 0.773 | 0.621 | 0.674 |
|  | 8 | 0.755 | 0.75 | 0.736 | 0.799 | 0.719 | 0.743 |
|  | 10 | 0.716 | 0.674 | 0.675 | 0.724 | 0.679 | 0.682 |
|  | Ensemble | 0.775 | 0.648 | 0.691 | 0.79 | 0.624 | 0.685 |
| MG | 4 | 0.459 | 0.264 | 0.343 | 0.44 | 0.249 | 0.325 |
|  | 6 | 0.443 | 0.276 | 0.344 | 0.471 | 0.264 | 0.347 |
|  | 8 | 0.438 | 0.292 | 0.351 | 0.473 | 0.289 | 0.364 |
|  | 10 | 0.481 | 0.281 | 0.362 | 0.463 | 0.272 | 0.349 |
|  | Ensemble | 0.526 | 0.261 | 0.365 | 0.537 | 0.254 | 0.364 |
| MN | 4 | 0.6 | 0.646 | 0.617 | 0.625 | 0.591 | 0.602 |
|  | 6 | 0.565 | 0.662 | 0.606 | 0.684 | 0.612 | 0.642 |
|  | 8 | 0.596 | 0.603 | 0.594 | 0.608 | 0.616 | 0.607 |
|  | 10 | 0.55 | 0.646 | 0.59 | 0.652 | 0.641 | 0.642 |
|  | Ensemble | 0.64 | 0.637 | 0.634 | 0.679 | 0.608 | 0.638 |
| ZN | 4 | 0.64 | 0.691 | 0.66 | 0.667 | 0.683 | 0.67 |
|  | 6 | 0.711 | 0.661 | 0.681 | 0.681 | 0.672 | 0.672 |
|  | 8 | 0.685 | 0.671 | 0.673 | 0.712 | 0.668 | 0.685 |
|  | 10 | 0.729 | 0.667 | 0.693 | 0.713 | 0.677 | 0.69 |
|  | Ensemble | 0.755 | 0.655 | 0.699 | 0.746 | 0.663 | 0.699 |

**Supplementary Table 3:**
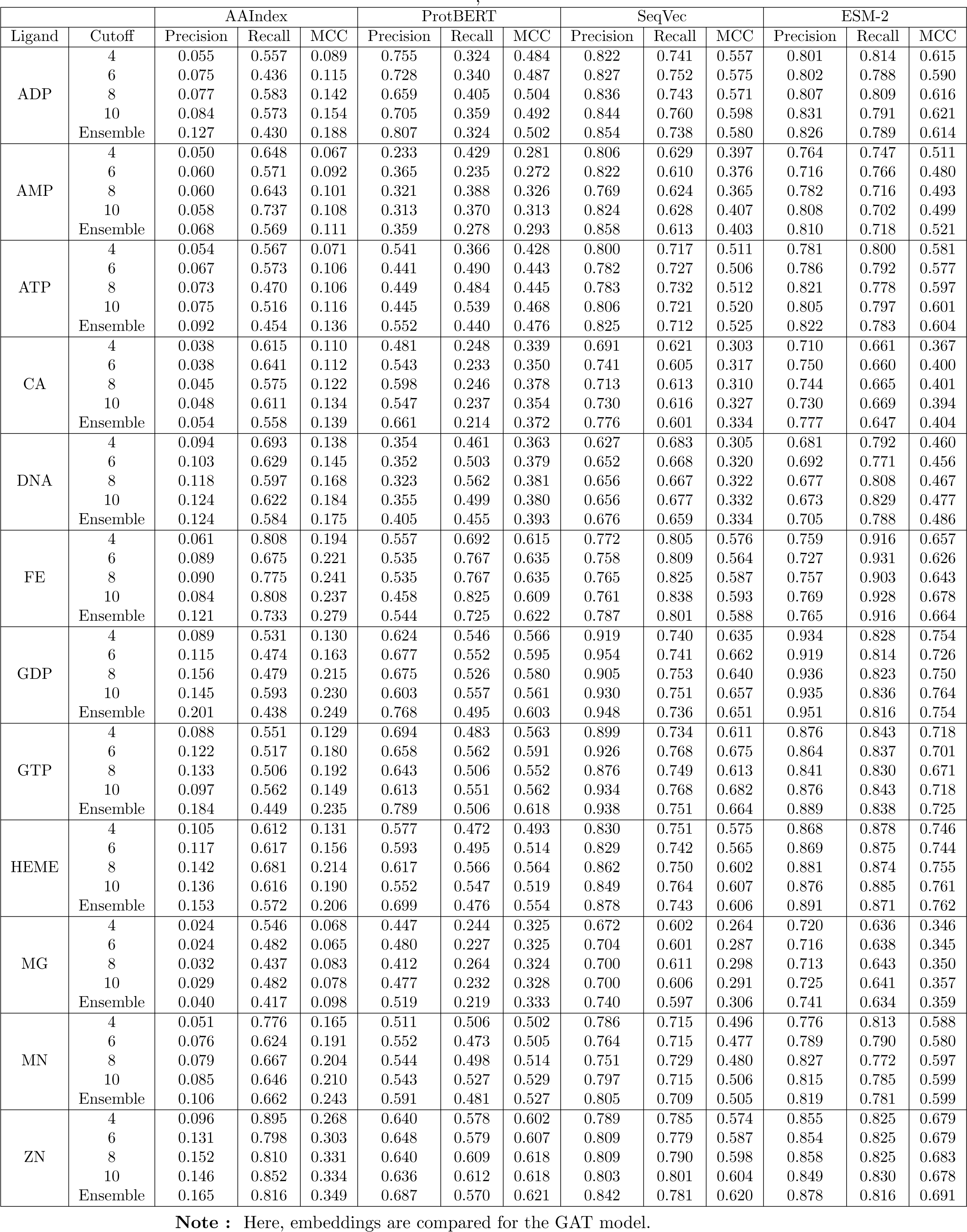
Comparison of different embeddings with different cutoff distances.

**Supplementary Table 4:**
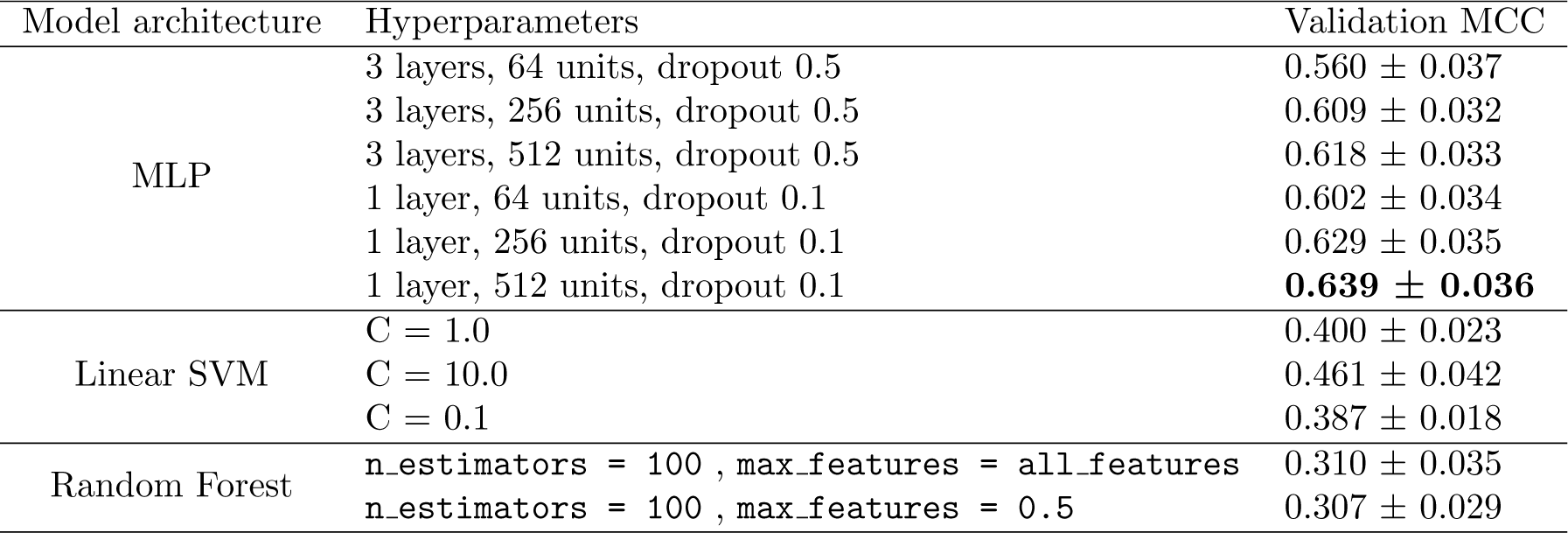
Models and hyperparameters used to select the sequence baseline. The displayed scores are means and standard deviations of validation MCC scores from 5-fold cross-validation on the ADP ligand training dataset.

**Supplementary Table 5:**
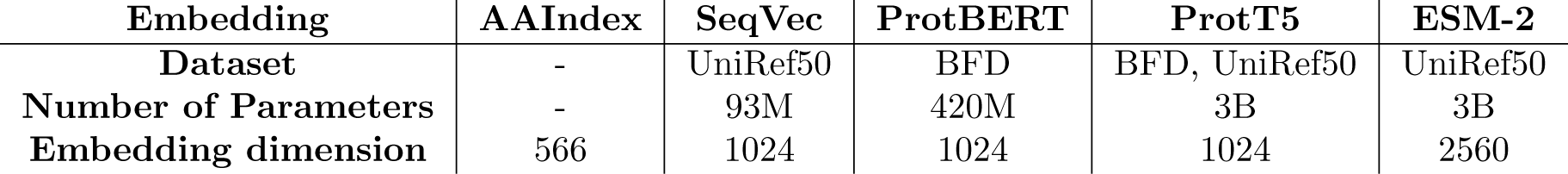
Comparison of embeddings.

| Embedding | AAIndex | SeqVec | ProtBERT | ProtT5 | ESM-2 |
| --- | --- | --- | --- | --- | --- |
| Dataset | - | UniRef50 | BFD | BFD, UniRef50 | UniRef50 |
| Number of Parameters | - | 93M | 420M | 3B | 3B |
| Embedding dimension | 566 | 1024 | 1024 | 1024 | 2560 |

**Supplementary Table 6:** Hyperparameter values tried in manual tuning.

| Hyperparameter | Values |
| --- | --- |
| Number of units in GNN layers | 64, 256, 512, 1024 |
| Learning rate | 3e-4, 1e-3 |
| Weight decay | 1e-5, 1e-2 |
| Dropout rate | 0, 0.3, 0.5 |
| Number of attention heads in GAT | 1, 2, 4 |
| Residual connections | True, False |
| Batch normalization | True, False |

**Supplementary Table 7:** Protein-DNA/RNA benchmarks summary. From GraphBind we used the protein-DNA benchmarking set consisting of a training set DNA Train 573 and a test set DNA Test 129, and we employed the protein-RNA benchmarking set consisting of a training set RNA Train 495 and a test set RNA Test 117. From GraphSite, we used the protein-DNA benchmarking test set DNA Test 181, and we trained the model on the same protein-DNA set DNA Train 573 from GraphBind. All protein-DNA/RNA benchmarks were downloaded in FASTA format, and underwent the same preprocessing strategy used for the Yu benchmark. We thus had to discard protein sequences with a high mismatch between the sequence from the benchmark and the sequence of residues from PDB.

| Dataset | Sequences | Binding residues | Non-Binding residues | Missing protein graphs |
| --- | --- | --- | --- | --- |
| DNA_Train_573 | 573 | 14479 | 145404 | 18 |
| DNA_Test_129 | 129 | 2240 | 35275 | 2 |
| DNA_Test_181 | 181 | 3208 | 72050 | 18 |
| RNA_Train_495 | 495 | 14609 | 122290 | 36 |
| RNA_Test_117 | 117 | 2031 | 35314 | 10 |

**Supplementary Table 8:** Comparison with existing methods - protein-DNA/RNA benchmarks.

| Dataset | Method | AUC | AUPR | MCC |
| --- | --- | --- | --- | --- |
| DNA_Test_129 | GraphBind | 0.928 | 0.519 | 0.499 |
|  | GraphSite | 0.919 | 0.502 | - |
|  | EquiPNAS | **0.943** | **0.582** | - |
|  | GeoBind | 0.940 | - | **0.526** |
|  | GAT8 + ProtT5 (ours) | 0.922 | 0.510 | 0.488 |
| DNA_Test_181 | GraphBind | 0.904 | 0.339 | 0.392 |
|  | GraphSite | 0.903 | 0.336 | **0.397** |
|  | EquiPNAS | **0.921** | **0.393** | - |
|  | GAT8 + ProtT5 (ours) | 0.898 | 0.337 | 0.364 |
| RNA_Test_117 | GraphBind | 0.854 | - | 0.322 |
|  | GeoBind | 0.874 | - | **0.373** |
|  | EquiPNAS | **0.887** | **0.320** |  |
|  | GAT8 + ProtT5 (ours) | 0.810 | 0.261 | 0.292 |

